# MENTOR: Multiplex Embedding of Networks for Team-Based Omics Research

**DOI:** 10.1101/2024.07.17.603821

**Authors:** Kyle A. Sullivan, J. Izaak Miller, Alice Townsend, Mallory Morgan, Matthew Lane, Mirko Pavicic, Manesh Shah, Mikaela Cashman, Daniel A. Jacobson

## Abstract

While the proliferation of data-driven omics technologies has continued to accelerate, methods of identifying relationships among large-scale changes from omics experiments have stagnated. It is therefore imperative to develop methods that can identify key mechanisms among one or more omics experiments in order to advance biological discovery. To solve this problem, here we describe the network-based algorithm MENTOR - Multiplex Embedding of Networks for Team-Based Omics Research. We demonstrate MENTOR’s utility as a supervised learning approach to successfully partition a gene set containing multiple ontological functions into their respective functions. Subsequently, we used MENTOR as an unsupervised learning approach to identify important biological functions pertaining to the host genetic architectures in *Populus trichocarpa* associated with microbial abundance of multiple taxa. Moreover, as open source software designed with scientific teams in mind, we demonstrate the ability to use the output of MENTOR to facilitate distributed interpretation of omics experiments.

## Introduction

As the amount of new omics data continues to proliferate, it is increasingly important to develop new technologies to integrate these data to understand complex biological systems. Network biology can be used to model biological entities (e.g., genes) as nodes, and the relationships or interactions among these genes can be represented as edges. Researchers can then combine multiple data layers into a single network structure and use network propagation methods to study known interactions or predict new ones. Network propagation can help identify relationships between biological molecules, such as proteins and metabolites ^1^.

Representing omics data as networks enables the discovery of novel interactions and relationships using network propagation and random walks ^2–4^. A random walk is a process in which a “walker” proceeds randomly from one node to an adjacent node^5^. The random walk method has been extended with a restart concept, forcing the walker to halt and return to the starting (“seed”) node^6,7^. This restart option effectively constrains the walk length without a hard threshold. This enhanced methodology is known as “Random Walk with Restart” (RWR). The result of running a random walker is a measure of the topological similarity of the seed node to all other nodes in the network, which can be used for clustering or other machine learning methods.

Valdeolivas et al. (2019) extended the RWR algorithm to multiplex networks. A multiplex network consists of multiple layers, each representing a different relationship between nodes^8^. For example, researchers may assemble a multiplex network from a co-expression network and a protein-protein interaction network. The multiplex network reduces the influence of spurious interactions, helping researchers identify high-confidence interactions for further investigation. In addition, the combination of multiple relationships into a single data structure for RWR improves gene-disease interaction prediction^4^. Multiplex networks can, therefore, leverage multiple types of biological evidence to identify relationships among a set of genes.

A common problem in biology is identifying relationships among a gene set. It is often unclear how genes implicated by one or more analyses are grouped biologically. For example, after analyzing one or more omics data types (e.g., differential gene expression by RNA-seq, genome-wide association study [GWAS], etc.), it is not always apparent how these genes functionally interact. Current tools to identify relationships among a gene set include Gene Ontology (GO^9^) enrichment for all omics types and weighted gene co-expression network analysis (WGCNA)^10^ for gene expression data. However, these methods’ limitations include that GO terms are manually curated and limited to known biological relationships, and is therefore not a comprehensive representation of all possible biological relationships. Moreover, WGCNA requires a complete expression dataset to identify gene models rather than identifying relationships among differentially expressed genes or a gene set of interest. Thus, it is essential to develop methods of identifying biological relationships among a gene set that can simultaneously focus on a bespoke set of genes while leveraging multiple independent types of biological evidence to establish these relationships.

To address this need, we created MENTOR (Multiplex Embedding of Networks For Team-Based Omics Research), a method for aiding mechanistic interpretation of omics datasets using RWR exploration of multiplex networks to determine the topological relationships among all genes in the set. As an abstraction of the edge density of these networks, a topological distance matrix is created and hierarchical clustering used to create a dendrogram representation of the functional interactions among the genes in the set. Thus, MENTOR enables users to find “clades” of functionally related genes using network analysis of multi-omics data. This work presents a software package enabling MENTOR as a computational systems biology workflow through a command-line interface.

## Methods

### Overview of MENTOR

MENTOR is a software extension to RWRtoolkit ^11^, which implements the random walk with restart (RWR) algorithm on multiplex networks using the RandomWalkRestartMH package in R^4^. Briefly, the RWR algorithm traverses a random walker across a monoplex or multiplex network using a single node, called the seed, as an initial starting point. All other genes within the network are assigned a score, representing the probability of the random walker reaching each gene starting from the seed. A random walk performed on a monoplex network only provides a single layer of biological evidence. In contrast, a multiplex network consists of multiple layers, each representing different lines of biological evidence. Multiplex networks have previously demonstrated a greater ability to recover biological interactions by maintaining the internal topology of each layer compared to combining all layers from each network^4^. A random walk on a multiplex network allows the walker to proceed from one layer to another via the cross-layer edges, and a restart probability prevents the walker from getting trapped within the topology of a single layer. The multiplex network also allows the topology of each individual layer to be preserved. MENTOR builds on these tools to enable users to find groups or clades of functionally related genes based on RWR embeddings. A multiplex network approach includes networks generated from multiple lines of evidence, such as gene expression data, protein-protein interaction data, and metabolomic interaction data, constituting millions of lines of evidence of complex biology. Because these lines of evidence are based on real-world experimental data, multiplex networks can leverage the unique network topology of thousands of experiments already performed by the biological research community to fully integrate multi-omics data and derive novel mechanistic connections.

An overview of the MENTOR workflow is shown in **Figure 1**. Starting from a single gene in the user’s gene set, a random walk exploration occurs from this gene, and all other genes in the multiplex network are given a score based on how frequently the genes are visited by the random walk. This process is then repeated for all other genes in the user’s gene set. After generating RWR-score vectors for each seed node, the vectors are sorted in rank order. The mean RWR score at each rank is then calculated [**Supplementary Fig. 1**]. For a given set of seed genes, the elbow^12^ of the mean scores-vs-ranks curve corresponds to a global rank at which the most closely associated nodes in the network have been found. The RWR vectors are then filtered, retaining only nodes that achieved a rank better than this maximum elbow rank for one of the seed nodes.

**Figure 1.**
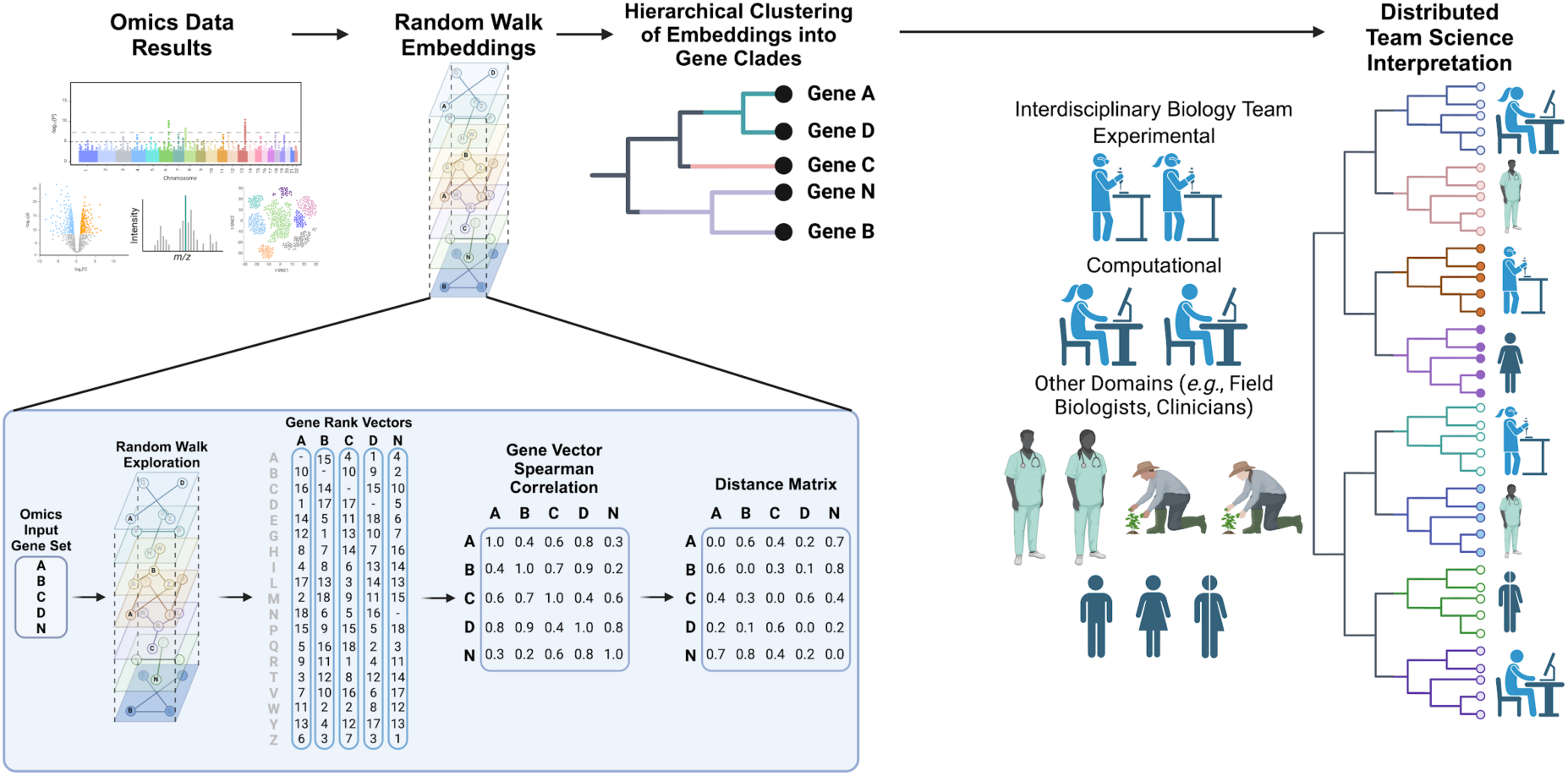
MENTOR Overview. Starting from a gene set derived from an omics dataset (e.g., GWAS, differential bulk or single-cell RNA-seq, differential protein abundance), users include a multiplex network for RWR exploration starting from each single gene in the set. This generates a rank-ordered vector of all genes explored in the multiplex when starting from this single gene. These rank vectors are then truncated to the most frequently explored genes (elbow point of RWR score), and rank vectors are compared by Spearman correlation. Euclidean distances of gene vectors are then calculated by 1 - *ρ*, with a distance of 0 indicating perfectly overlapping vectors and a distance of 1 illustrating completely dissimilar vectors. These Euclidean distances, representing multiplex embeddings of the networks from each individual gene, are then arranged into a dendrogram using agglomerative hierarchical clustering. The dendrogram is then cut at a distance threshold sufficient to divide the dendrogram into a discrete number of clades. Finally, clades are then distributed for interpretation, with each clade containing genes that are closely interconnected in the multiplex network based on RWR-derived embeddings. Figure made with Biorender.com.

### Clustering

Each node’s RWR vector represents that node in the context of the multiplex network. Clustering the feature vectors thus identifies the most similar nodes based on network topology. The rank order similarity of the feature vectors is calculated using Spearman’s rank correlation coefficient *ρ*. The correlations are then converted to distances (*1* − *ρ*). Based on the Spearman distances, the genes are clustered using agglomerative hierarchical clustering in the R stats package^13^. The *cutree* function from the R stats package is then used, with a specified *k,* to group genes into the desired number of gene groups, referred to as clades. Iterative sub-clustering can be performed on the dendrogram visualization to ensure that the number of genes within each clade does not exceed a specified maximum size. The process begins with an initial number of clusters using the *cutree* function to split the dendrogram. The size of each cluster is then assessed. If the cluster size is smaller than the maximum allowed, the cluster labels are saved, and no further action is taken for that cluster. Conversely, if any cluster exceeds the maximum size, the number of clusters in the *cutree* function is increased by a user-defined amount. The sizes of the remaining clusters are evaluated again. If they are within the maximum size, their labels are saved. This process is repeated until no clusters exceed the maximum size.

### Outputs

A summary of inputs and outputs to MENTOR is listed in **Supplementary Figure 2**. MENTOR provides the user with three primary outputs: (i) clade labels for the input genes, (ii) a dissimilarity matrix, and (iii) a dendrogram representation of the hierarchical clustering. By default, cluster labels are assigned using hierarchical clustering of the Spearman distances, with a default specification of *k* equal to three clades, and an associated dendrogram visualization is constructed using the clustering results. The user also has the option to explore the dendrogram and select different clusters by visual inspection, adjust the parameter *k* to the desired number of clades, and run sub-clustering to ensure the number of genes within each clade does not exceed a maximum size. The user can then pass the gene clusters to further downstream analysis methods such as GO enrichment^9^.

### Additional MENTOR options

A mapping file can be provided by the user to adjust the labels of the dendrogram leaves. For example, Ensembl IDs can be mapped to approved gene symbol labels. A heatmap file can also be provided that represents the data sources from which a given gene originated. For example, in the case of single-cell RNA-seq data, the rows of the heatmap file can represent the different cell types within which a gene is either up- or down-regulated (e.g., log_2_ fold change). If the user provides a heatmap file, then a heatmap visualization is affixed to the right side of the dendrogram visualization, which allows the user to discern both horizontal (across cell type) and vertical (cell-type-specific) patterns of gene regulation (**Supplementary Figure 2C)**. For instance, within single-cell RNA-seq data this can allow the user to quickly discern patterns of up- and down-regulation both shared and unique to the different cell types within the data source. In the case of other data sources, such as GWAS results, the gene is either implicated or not and color is used to represent either presence or absence of an association and the magnitude and direction of effect size. When the number of genes within the user’s input gene list increases significantly, which is particularly relevant to single-cell data, an additional argument can be provided which specifies for the dendrogram to be plotted within polar coordinates. This not only reduces the overall size of the dendrogram but allows for faster interpretation of broad functional changes across data sources. All additional options are presented in the GitHub repository with detailed descriptions that adjust various aspects of the visualizations such as the plot dimensions and custom color assignments.

### Human Phenotype Ontology (HPO) gene set clustering

We tested MENTOR’s capability to partition genes into functional groups in a supervised manner using gene sets derived from the Human Phenotype Ontology (HPO^14^). First, we combined all genes from two different HPO terms and labeled each gold-standard gene with the associated HPO term, and then used MENTOR to cluster these genes based on multiplex RWR exploration. HPO is a good representation of a typical use case that a biologist might have (e.g., from a GWAS or differential expression analysis), as each HPO term is associated with a set of genes associated with a phenotype. After MENTOR clustered all genes from a pair of HPO terms, the clustering was evaluated by using the dissimilarity matrix to calculate pairwise distances of each gene within each HPO term and between HPO terms. This generated the mean pairwise within-term and between-cluster distances, where a high quality clustering should have low within-cluster distances and high between-cluster distances.

We used three pairs of HPO terms for testing MENTOR. This included “abnormal circulating glycine concentration” (HP:0010895) and “left-to-right shunt” (HP:0012382); “abnormal circulating glycine concentration” (HP:0010895) and “limited neck range of motion” (HP:0000466); and “abnormality of the protein C anticoagulant pathway” (HP:0030780) and “primary peritoneal carcinoma” (HP:0030406). All MENTOR clusters were generated using a multiplex network we created from the eight network layers of the HumanNet V3 network ^15^ (**Table 1**).

**Table 1.** Summary of HumanNet V3 network layers in multiplex network. The number of nodes in the multiplex is the union of the nodes in the layers. The number of edges in the multiplex is the sum of the edges in each layer.

| Network Description | No. Nodes | No. Edges |
| --- | --- | --- |
| Co-citation | 18,206 | 1,073,857 |
| Co-expression | 12,152 | 80,382 |
| Gene neighborhood | 2,329 | 97,209 |
| Genetic interaction | 10,453 | 174,068 |
| Pathway database | 8,515 | 134,312 |
| Phylogenetic profile associations | 2,123 | 16,454 |
| Protein domain profile associations | 12,618 | 72,729 |
| Protein-protein interaction network | 17,778 | 630,893 |
| Multiplex | 18,488 | 2,279,904 |

**Table 2.** Summary of *P. trichocarpa* network layers and multiplex. Networks are derived from *P. trichocarpa* data unless otherwise noted. The number of nodes in the multiplex is the union of the nodes in the layers. The number of edges in the multiplex is the sum of the edges in each layer.

| Network Description | No. Genes | No. Edges | Citation |
| --- | --- | --- | --- |
| ATRM (*A. thaliana* orthologs) | 749 | 1,687 | Jin et al. (2015)<sup>82</sup> |
| BWERF | 2,134 | 11,694 | Newly generated |
| Co-expression atlas | 4,253 | 32,547 | J. Zhang et al. (2018)<sup>83</sup> |
| Co-knockout (*A. thaliana* orthologs) | 2,422 | 159,006 | Oellrich et al. (2015)<sup>84</sup> |
| Co-metabolite | 4,108 | 46,175 | P. Zhang et al. (2010)<sup>85</sup> |
| PlantRegMap | 16,688 | 90,812 | Jin et al. (2015)<sup>82</sup>; Jin et al. (2017)<sup>86</sup>; Tian et al. (2020)<sup>87</sup> |
| STRING-DB PPI (*A. thaliana* orthologs) | 14,133 | 218,083 | Szklarczyk et al. (2021)<sup>88</sup> |
| iRF-predicted expression (leaf) | 15,204 | 209,824 | Cliff et al. (2019)<sup>89</sup> |
| iRF-predicted expression (xylem) | 38,872 | 138,194 | Cliff et al. (2019)<sup>89</sup> |
| Multiplex | 41,764 | 908,022 |  |

### *Populus trichocarpa* Microbial Colonization GWAS

RNA-seq data from xylem tissues of a diverse population of *P. trichocarpa* genotypes were obtained and reads were aligned, trimmed, and filtered against the *P. trichocarpa* v3.0 reference genome from Phytozome^16–18^. Reads that did not map to the reference genome were considered putative microbiome members and included in subsequent metatranscriptomic analyses. These unmapped reads were taxonomically classified at the genus level using ParaKraken^19^. Only taxa with a pseudo-abundance of at least 1% were retained, and sample-level normalization was applied to address sequencing biases. Outlier taxa were identified and excluded from the dataset. The finalized set of microbial taxa was then binarized, representing presence or absence, and utilized as phenotypic data for GWAS (**Supplementary Table 1**).

Ascomycetes genera *Stagonosporopsis* and *Ilyonectria* were selected for further analysis due to their identification as known tree-root fungi in *Populus deltoides* and *P. trichocarpa* respectively. Both genera function as root endophytes but under certain conditions can demonstrate pathogenicity in roots or vasculature^20–22^. We ran a GWAS on the binarized phenotypes for both *Stagonosporopsis* and *Ilyonectria* with 6M *P. trichocarpa* v3.0 single nucleotide polymorphisms (SNPs). SNPs were filtered for quality, missing genotypes (--geno 0.15) missing individuals (--mind 0.15), Hardy-Weinburg equilibrium (--hwe 1E-50), for a minor allele frequency threshold of 0.05 using PLINK v1.9^23–25^. GAPIT version 3^26^ was used to perform GWAS using MLM^27^, MLMM^28^, FarmCPU^29^, and BLINK^30^ models. Significant SNPs (FDR-adjusted p-value ≤ 0.1) across all models for both *Stagonospora* and *Ilyonectria* were assigned to the closest gene using a modified version of bedtools --closest function^31^ and the poplar v3.1 .gff file obtained from Phytozome^17^. We combined the genes identified across all models for each fungal taxa as the input gene set and used a previously described *P*. *trichocarpa* multiplex^18^ as input to MENTOR. The genes in the dendrogram output were divided across team members for interpretation in the context of host-microbial interactions.

## Results

### MENTOR separates genes associated with different Human Phenotype Ontology terms based on network topology

In order to test whether MENTOR could separate distinct biological processes into separate clades, we applied this method to a single input composed of genes from three pairs of Human Phenotype Ontology (HPO) terms. From the dissimilarity matrix computed by MENTOR, we computed the pairwise similarity of all pairs of genes within a single HPO term, as well as the similarity between the genes from the two different HPO terms. In each of the three exemplar gene sets, MENTOR partitioned the genes using network topology, with clades containing genes which largely constituted a single HPO label (**Figure 2**). One of the genes labeled as “abnormal circulating glycine concentration” did not cluster with the other genes with this label when combining with genes associated with “left-to-right shunt”, indicating it may have a distinct function from the others (**Figure 2A**). Similarly, all but one gene labeled as “abnormal circulating glycine concentration” were in the same clade when combined with genes associated with “limited neck range movement” (**Figure 2B**). Finally, genes labeled as “abnormal protein C anticoagulant pathway” and “primary peritoneal carcinoma” clustered perfectly together in clades based on network topology (**Figure 2C**).

**Figure 2.**
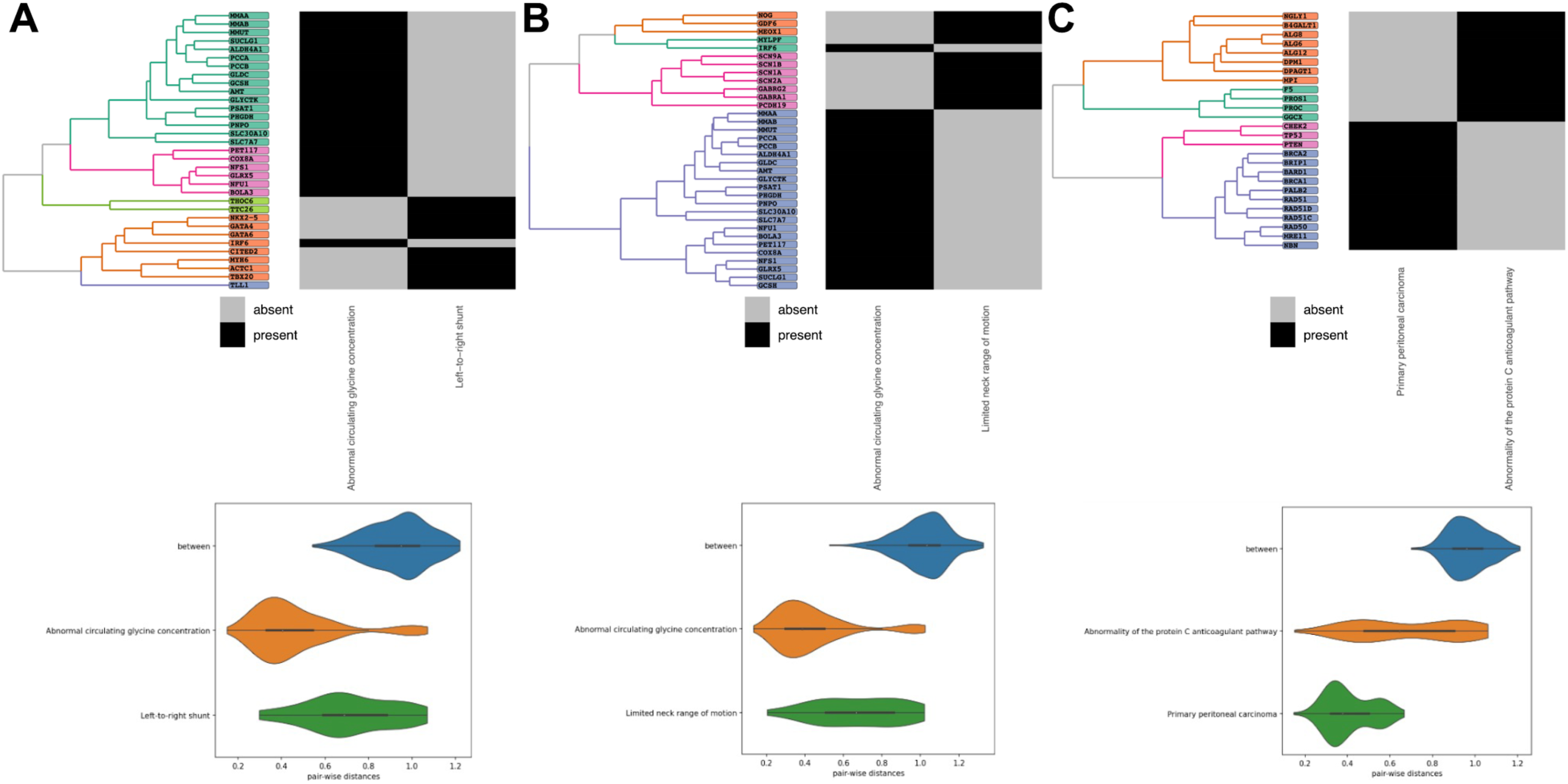
MENTOR separates genes associated with distinct phenotypes based on network embeddings. When mixing two HPO terms into an input gene list, MENTOR largely separates these genes into distinct clades. **A)** Mixing HPO terms for “abnormal circulating glycine concentration” and “left-to-right shunt” separates genes into five clades clustered by each HPO term with the exception of only a single intermixed gene. This corresponded to a high mean pairwise distance of genes between HPO terms with low within-term distances. This pattern was also consistent when combining **B)** “abnormal circulating glycine concentration” and “limited neck range of motion” as well as **C)** “abnormality of the protein C anticoagulant pathway” and “primary peritoneal carcinoma” HPO terms.

### MENTOR identifies functional relationships among *P. trichocarpa* genes associated with fungal colonization

Next, we performed a genome-wide association study (GWAS) of microbial colonization in *Populus trichocarpa* to demonstrate an example MENTOR pipeline and its functionality (**Figure 3, Supplementary Table 1**). The *Stagonosporopsis* GWAS identified 171 unique significant SNPs (FDR-adjusted p-value ≤ 0.1) and 32 unique genes combined across the four models. The *Ilyonectria* GWAS identified 212 unique significant SNPs (FDR-adjusted p-value ≤ 0.1) and 42 unique genes combined across the four models. Two different SNPs from the *Stagonosporopsis* and *Ilyonectria* GWASs were mapped to the same gene, Potri.002G232500. In the *Stagonosporopsis* GWAS, SNP Chr02_22504473 was identified by the MLM model (FDR-adjusted p-value = 7.70E-02) and was found 7265 base pairs (bp) upstream of Potri.002G232500. In the *Ilyonectria* GWAS, SNP Chr02_22498716 was identified by both the FarmCPU (FDR-adjusted p-value =8.85E-02) and BLINK (FDR-adjusted p-value = 2.96E-02) models and located 1508 bp upstream of Potri.002G232500.

**Figure 3.**
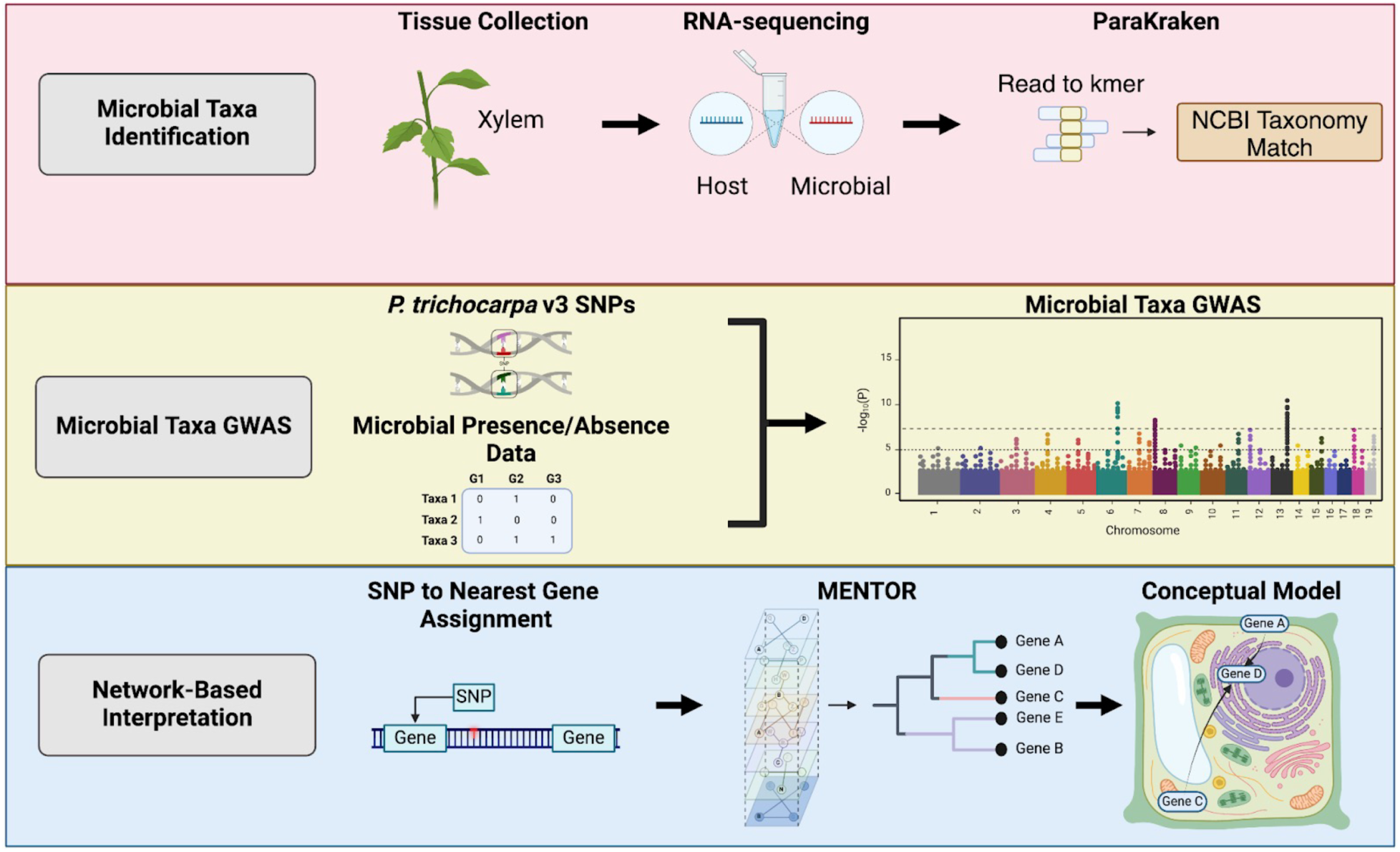
Workflow for MENTOR interpretation of *Populus trichocarpa* microbial taxa GWAS. Figure made with Biorender.com.

All 73 unique genes (*Stagonosporopsis* = 32, *Ilyonectria* = 42, 1 shared) in the input gene set were found in the *P. trichocarpa* multiplex used by MENTOR. In total, 8 clusters of mechanistically related genes were generated and can be visualized by the MENTOR output dendrogram (**Figure 4A**). Interpretation of the MENTOR dendrogram revealed population-level variation in genes encoding proteins involved in microbial colonization-related activities in *P. trichocarpa*; particularly in cell-surface receptors, modulation of fungal growth, cell wall integrity sensing, Ca^2+^ influx and reactive oxygen species (ROS) signaling, and transcriptional reprogramming of growth and immunity tradeoffs (**Figure 4B**).

**Figure 4.**
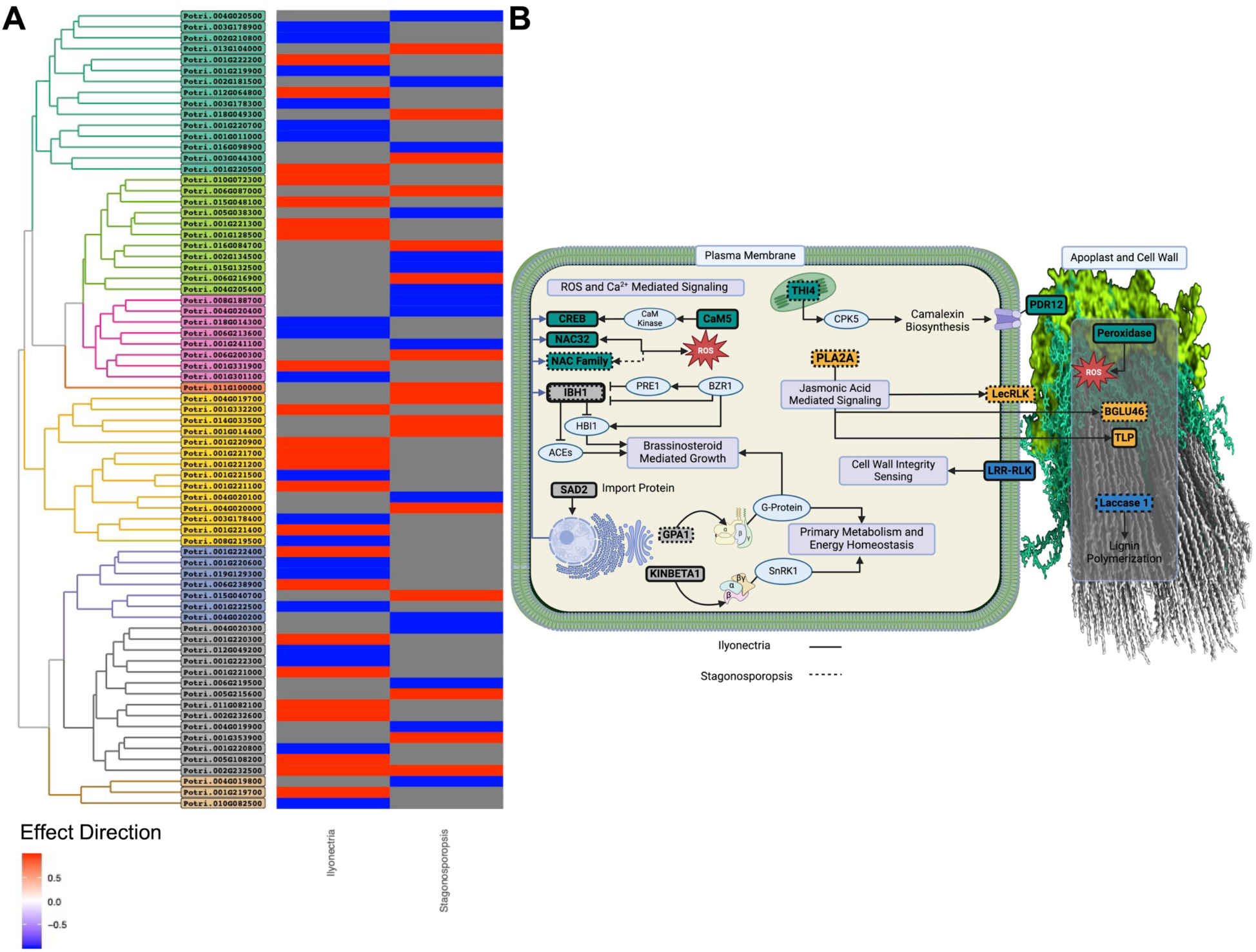
MENTOR identifies key biological pathways related to host genes involved in *Populus trichocarpa* microbial taxa abundance. **A)** MENTOR dendrogram of *Populus trichocarpa* host genes associated with colonization (presence/absence) of *Ilynocetria* and *Stagonosporopsis* taxa. Heatmap indicates direction of effect of SNP associated with nearest gene. **B)** Conceptual model of genes associated with fungal taxa color-coded by each MENTOR clade, including secondary cell wall visualization identified from Addison et al., 2024. Figure made with Biorender.com.

We identified two genes that encode transmembrane cell surface receptor proteins in distinct clades likely involved in the recognition of each *Stagonosporopsis* and *Ilyonectria* taxa. SNP Chr01_992890 was identified by FarmCPU (FDR-adjusted p-value = 2.88E-02) in the *Stagonosporopsis* GWAS and is found in the gene body of Potri.001G014400, which was clustered into the yellow clade and encodes a G-type lectin receptor-like kinase (LecRLK). LecRLKs are cell surface receptors that are widely recognized to confer resistance via their roles in pathogen recognition and immune response^32,33^ as well as their roles in defense signaling^34^. Similarly, SNP Chr19_15551698 was identified by the FarmCPU model (FDR-adjusted p-value = 9.75E-03) in the *Ilyonectria* GWAS and is found in the gene body of Potri.019G129300, which was clustered into the blue clade and encodes a leucine-rich repeat receptor-like kinase (LRR-RLK). LRR-RLKs are also cell surface receptors that have important roles in microbial recognition and plant immune responses, particularly in the recognition of pathogen associated molecular patterns (PAMPs) to activate downstream signaling pathways^35^.

In addition to LecRLKs widespread role in fungal detection, some LecRLKs are regulated by essential JA signaling components^34^. Consistent with this, we found population-level variation in genes related to fine-tuning the modulation of fungal growth through jasmonic acid (JA) signaling. Li et al. (2004)^36^ demonstrated that the *A. thaliana* ortholog of the LecRLK we identified (AT5G60900) was significantly upregulated in transgenic plants overexpressing WRKY70, a gene involved in JA signaling that enhances plant defense against pathogens. In addition, we identified SNPs in the *Ilyonectria* GWAS that mapped to genes encoding multiple types of pathogenesis-related (PR) proteins. Specifically, the MLM revealed numerous SNPs mapping to membrane localized thaumatin-like proteins (PR-5s). PR-5s are induced by abiotic and biotic stressors and modulate fungal invasion through direct antifungal effects in plants^37,38^. They also act as elicitors of additional antifungal proteins in *Populus* spp. that contribute to increased disease resistance^39^. PR-5s in rice and in wheat are also induced in response to JA signaling^40,41^, highlighting their central roles in the intersection of antifungal activity and JA response.

The MLM model in the *Stagonosporopsis* GWAS also identified SNP Chr04_1307502 (FDR-adjusted p-value = 6.34E-02) and mapped to Potri.004G019700, which encodes a β-glucosidase. The *A. thaliana* ortholog was β-glucosidase 46 (BGLU46), which is localized in the extracellular region and involved in hydrolysing monolignol glucosides during lignification^42^ (Escamilla-Treviño et al., 2006), an essential process for cell wall integrity and to resist fungal invasion^43^. In addition, β-glucosidases from this family can also be induced by JA in phytohormone signaling and involved in the activation of chemical defense compounds that enhance resistance^44^.

In the *Stagonosporopsis* GWAS, the MLM model identified two significant SNPs (Chr14_2748605, FDR-adjusted p-value = 4.28E-02; Chr14_2749599, FDR-adjusted p-value = 1.98E-02) found in the gene body of Potri.014G033500, which encodes a lipid hydrolysis protein phospholipase A2A (PLA2A). In *A. thaliana*, PLA2A is strongly induced by fungal infection and dependent on JA signaling; however, overexpression of PLA2A is linked to increased susceptibility, as pathogens exploit the lipolytic activity of this protein to facilitate host colonization^45^.

As cell surface receptors, LRR-RLKs integrate diverse microbial detection signals into downstream immune and symbiotic responses^46^. In particular, we found population-level variation in genes related to cell wall integrity sensing and cell wall changes associated with microbial-induced lignification. The *A. thaliana* ortholog to the LRR-RLK identified by the *Ilyonectria* GWAS (Potri.019G129300) is MALE DISCOVERER 1-INTERACTING RECEPTOR LIKE KINASE 2 (MIK2), which is required for resistance against fungal root pathogen *Fusarium oxysporum*; loss-of-function mutations in MIK2 result in compromised immune responses, characterized by reduced expression of immune marker genes, decreased jasmonic acid production, and impaired lignin deposition^47^. Additionally, the MLM model in the *Stagonosporopsis* GWAS identified multiple significant SNPs mapping to Potri.015G040700, which is orthologous to LACCASE 1 (LAC1) in *A. thaliana*. Extracellular laccases play crucial roles in the lignification of secondary cell walls, a fundamental process that not only contributes to the structural integrity and development of the cell but also serves as both a constitutive and inducible response to pathogen defense^48–50^. For example, in *Gossypium hirsutum* (upland cotton), overexpression of a lignin-associated laccase (GhLAC15) enhanced resistance to Verticillium wilt by increasing total lignin content and altering lignin structure and other cell wall traits^49^.

Ca^2+^ influx and ROS signaling are some of the earliest responses detected in plant cells upon microbial sensing and invasion and are heavily involved in plant immunity^51^. We found population-level variation in cellular responses linking Ca^2+^ influx and reactive oxygen species (ROS) signaling cascades to antimicrobial compound production and secretion. In the *Stagonosporopsis* GWAS, we identified multiple SNPs from the MLM model mapping to the gene Potri.004G020500 encoding *THIAZOLE SYNTHASE 1* (THI1), a critical enzyme in thiamine (vitamin B1) biosynthesis. Thiamine functions to prime plant defenses, activating systemic acquired resistance (SAR) in plants to provide extended and efficient resistance to a broad spectrum of bacterial, fungal, and viral pathogens^52^. Additionally, THI1 interacts directly with calcium-dependent protein kinase 5 (CPK5) in *A. thaliana*^53^, and CPK5 acts as a hub within Ca^2+^ signaling to induce SAR^54^. Moreover, CPK5 regulates the production of the most prominent phytoalexin, camalexin, in *A. thaliana*, an antimicrobial compound produced by pathogen invasion that enhances disease resistance^55^. The MLM model in the *Ilyonectria* GWAS identified SNP Chr03_18668538 (FDR-adjusted p-value = 2.90E-02) mapping to gene Potri.003G178900, which encodes *PLEIOTROPIC DRUG RESISTANCE 12* (PDR12), an ATP-binding cassette (ABC) transporter located in the plasma membrane in *A. thaliana* found to regulate the release of camalexin in response to necrotrophic fungus *Botrytis cinerea* infection^56^.

We found additional evidence integrating other genes in the Ca^2+^ signaling cascade. Specifically, multiple SNPs identified by MLM, MLMM, and FarmCPU models in the *Ilyonectria* GWAS were mapped within the gene body of Potri.001G222200, which encodes a calmodulin (CaM) 5 protein. CaMs bind to Ca^2+^ and can act as both signal receptors and transducers, depending on the isoform, and activate plant disease resistance responses^57^. In addition, SNP Chr01_22642160 was identified by the MLM model (FDR-adjusted p-value = 2.41E-02) and mapped to Potri.001G220700, a gene encoding adenosine 3’,5’-monophosphate (cAMP) response element-binding protein (CREB). CREB proteins are transcription factors that integrate Ca^2+^ signals into transcriptional responses via phosphorylation by CaM-dependent kinases, which are activated by CaMs altered through Ca^2+^ binding^58^.

Extracellular peroxidases are also heavily involved in the formation of hydrogen peroxides, specifically as part of ROS production leading to the oxidative burst response, which is a key early response to microbial colonization^59^. In the *Ilyonectria* GWAS, SNP Chr01_693267 (FDR-adjusted p-value = 1.46E-07) was identified by FarmCPU model was mapped to Potri.001G011000, which encodes a peroxidase. Bindschedler et al. (2006)^60^ found cell-wall localized peroxidases in *A. thaliana* play a crucial role in generating ROS in response to *F. oxysporum*; transgenic plants exhibiting which suppressed transcription of these peroxidases were more susceptible to both fungal and bacterial pathogens. Additionally, NO APICAL MERISTEM/*ARABIDOPSIS* TRANSCRIPTION ACTIVATION FACTOR/CUP-SHAPED COTYLEDON (NAC) transcription factors are also recognized as regulators of oxidative stress^61^. The MLM models in both the *Ilyonectria* and *Stagonosporopsis* GWASs identified SNPs mapping to Potri.001G220500 and Potri.018G049300, respectively, both of which encode NAC transcription factors. The *A. thaliana* ortholog of Potri.001G220500 is NAC032, which was identified to aid in detoxification within an oxidative stress regulatory network that includes additional NAC transcription factors^62^. NAC032 was also found to be co-expressed with plant glycosyltransferases that help regulate cellular redox status and detoxify ROS-reactive secondary metabolites in response to *Pseudomonas syringae* pv. *tomato* infection^63^.

We found population-level variation in genes involved in transcriptional reprogramming of brassinosteroid (BR)-mediated growth and primary metabolic activities that may influence growth-immunity tradeoffs. Both the *Stagonosporopsis* and *Ilyonectria* GWASs identified Potri.002G23250 in the gray clade, which was annotated as *INCLINATION1 BINDING BHLH PROTEIN1* (IBH1). IBH1 is an atypical basic helix loop helix (bHLH) transcription factor that negatively regulates brassinosteroid signaling^64^. Specifically, *BRASSINAZOLE RESISTANT 1* (BZR1) is a master transcriptional regulator in *Arabidopsis thaliana* that meditates growth and immunity tradeoffs^65^ and physically interacts with and inhibits IBH1 in the presence of BR^66^. Further, IBH1 is involved in antagonistic relationships with many BR-mediated growth promoting transcription factors, including *PACLOBUTRAZOL RESISTANCE 1* (PRE1), *ACTIVATORS FOR CELL ELONGATION* (ACE1-3), and *HOMOLOG OF BRASSINOSTEROID ENHANCED EXPRESSION2 INTERACTING WITH IBH1* (HBI1). Specifically, PRE1 and HBI1 are first activated by BZR1; PRE1 subsequently binds to IBH1 to prevent its inhibitory activity of ACEs and HBI to promote cell elongation and growth^66,67^. HBI1 is also a demonstrated negative regulator of pathogen-associated molecular pattern (PAMP) triggered immunity (PTI), where overexpression of HBI1 diminishes plant immune responses and can increase plant susceptibility to pathogens^68^. Additionally, both HBI1 and BZR1 have exhibited a trade-off between growth and immunity^65,69^. Therefore, interactions between IBH1 and these other transcription factors in the BR-mediated pathway likely modulate both the manner in which and how effectively plants respond to pathogen attacks.

The MLM model identified SNP Chr02_22516720 in the *Ilyonectria* GWAS that mapped to Potri.002G232600, which encodes an importin β. Importin β is a highly conserved karyopherin involved in chaperoning and transporting protein complexes, such as transcription factors, from the cytoplasm into the nucleus^70,71^. The *A. thaliana* ortholog of Potri.002G232600 is SUPER SENSITIVE TO ABA AND DROUGHT 2 (SAD2), which showed enhanced resistance to *P. syringae* pv. Tomato in overexpressing lines and susceptibility in knockout mutants due to the differences in expression of key defense response genes, such as EFR, MYB, and bHLH transcription factors regulated by SAD2^72^. With this model, we hypothesize that Potri.002G232600 may also regulate the expression of the previously mentioned bHLH transcription factors though its chaperone and transport activities and therefore contribute to additional growth-immunity tradeoffs during microbial colonization.

Both the FarmCPU and MLM models in the *Stagonosporopsis* GWAS identified SNP Chr06_23189402 which mapped to Potri.006G219500. The *A. thaliana* ortholog of Potri.006G219500 is *G PROTEIN ALPHA SUBUNIT 1* (GPA1), which encodes the alpha subunit of heterotrimeric G-proteins. G-proteins are signal mediators in developmental signal transduction and stress responses, including roles as a positive regulator in cell division and as a regulator of cell wall composition^73^. Loss of function of the Gα subunit (encoded by GPA1) results in enhanced resistance against a necrotrophic fungal pathogen^74^, suggesting a G-proteins may also contribute to a tradeoff between growth and immunity in response to microbial colonization. Heterotrimeric sucrose non-fermenting 1 (SNF1)/SNF1-related kinase 1 (SnRK1) protein kinases also integrate cellular responses to nutrient, energy, and stress signals to maintain cellular energy homeostasis^75^. The MLM model in the *Ilyonectria* GWAS identified multiple SNPs mapping to Potri.001G220800, which encodes the beta subunit of SnRK1 (KINBETA1 in *A. thaliana*). In addition to maintaining cellular energy homoeostasis, SnRK1 has been implicated in transcriptional reprogramming to balance growth-immunity tradeoffs and has essential roles in plant-pathogen interactions^76,77^. In rice, for example, overexpression lines containing a gene encoding the beta subunit of SnRK1 exhibited enhanced resistance against the fungal pathogen *Magnaporthe oryzae* and the bacterial pathogen *Xanthomonas oryzae* pv. *oryzae* (*Xoo*), whereas knockout lines were more susceptible to both pathogens^77^.

## Discussion

MENTOR is a computational systems biology workflow and software package that conveniently clusters genes by functional interaction. MENTOR is designed for data integration and fusion, addressing the complexities of merging samples from different populations and data types where traditional AI or correlation methods fall short. For example, MENTOR can leverage differentially expressed genes from bulk and single-cell RNA-seq results, differentially abundant proteins from proteomics results, and significant genetic variants associated with a disease from GWAS results. Depending on the scientific question of interest, the union, intersection, or a specific combination of the results from these data sources can be fed into MENTOR as a single input gene list.

MENTOR uses hierarchical clustering to identify clades of functionally interacting genes (as represented by multiplex network topology), which can be intuitively visualized as a dendrogram. Common gene discovery and identification approaches such as GWAS and RNA-sequencing experiments generate hundreds or even thousands of randomly ordered genes, leaving researchers with difficult decisions on which genes to prioritize and explore further. MENTOR’s visualization enables team-based science to accelerate discovery by reducing the overwhelming number of genes into clades that can be divided across teams for interpretation (**Figure 1**). Moreover, the visualization’s hierarchical structure of the gene set helps orient researchers towards a mechanistic interpretation of clades of interest, as functionally interacting genes are clustered together. This collaborative framework enables parallelized interpretations to be unified into a single comprehensive mechanistic conceptual model, combining multiple perspectives into a cohesive narrative, thereby reducing the redundancy of having multiple individual reports. Taken together, MENTOR enhances previous methods, allowing the grouping of genes that are functionally interacting with one another, which significantly advances our ability to interpret complex biological datasets.

MENTOR is distinct from other hierarchical clustering methods, such as WGCNA^10^. Both MENTOR and WGCNA use hierarchical clustering to generate a dendrogram and “clades” or “modules” of genes, but they are implemented with different underlying purposes. In WGCNA, a dissimilarity matrix is used with Pearson correlation to create a gene coexpression network from the expression values associated with the input gene set and then performs hierarchical clustering to form modules. MENTOR uses hierarchical clustering to create a dendrogram from the distance metrics resulting from random walk with restart across a multiplex network, which is generated independently from the input gene set and is a required input for MENTOR, comprising known biological relationships relevant to the gene sets to explore. Notably, these lines of evidence are independent of the user’s dataset. Importantly, MENTOR does not cluster the full dataset itself from an omics experiment, but instead performs clustering based on the interconnected gene-gene relationships that input genes have with other genes in the multiplex network. These relationships are not based on phylogeny or sequence similarity, but rather by a network topological association determined by random walk with restart starting from the input gene set, traversing the network, and generating a distance matrix derived from rank order vectors for genes most highly connected to those within the input gene set. Here, we are effectively leveraging extensive knowledge from real-world experiments that we can use to support or refute hypotheses of mechanistic interaction, which allows us to identify connectivity between genes based upon real-world experimental data.

We first tested the extent to which MENTOR could use embeddings of network topology to separate genes associated with distinct biological processes into separate clades. We demonstrated three pairs of HPO terms largely clustered into clades containing genes from the same HPO term. We observed that when two HPO terms had low average values of pairwise semantic similarity (corresponding to a higher pairwise distance), the genes were partitioned into largely distinct clades corresponding to their associated HPO term. For example, when combining genes from the “abnormality of the protein C anticoagulant pathway” term with genes in the “primary peritoneal carcinoma” term, all carcinoma genes were contained within a single clade and all protein C anticoagulant pathway genes were partitioned into a separate clade. However, there were notable distinctions where genes from one HPO term instead clustered with genes corresponding to the alternate HPO term, *e.g.*, *IRF6* clustering with “abnormal circulating glycine concentration” instead of “Left-to-right shunt or “limited neck range of motion”). However, a study by Wu et al.^78^ explored the interaction between *IRF6* and glycine receptor beta, demonstrating a protective effect against the development of nonsyndromic cleft lip with or without cleft palate in the Han Chinese population. This study identified specific single-nucleotide polymorphisms (SNPs) in IRF6 that showed protective effects against the condition, indicating a potential role for IRF6 in glycine-related pathways. Thus, it is possible that this result may be explained by pleiotropy, as a gene may have multiple biological functions that are yet uncaptured in HPO or GO. There are notable limitations to biological ontologies^79–81^, including the fact that they are often curated manually based on individual biological expertise. This means that while the information contained within HPO and GO is likely reliable (few false positive gene-phenotype associations), there are many associations yet to be incorporated (*i.e.*, many false negative gene-phenotype associations). Therefore, a gene may, in fact, contribute to multiple processes across multiple clades in MENTOR, and the visualized grouping is based on the shortest possible clustering based on RWR vector comparisons combined with hierarchical clustering.

Finally, we demonstrated that MENTOR can be applied to GWAS data in order to identify functional relationships among these genes. After identifying genetic variants associated with the colonization of two fungal taxa, we employed MENTOR to contextualize host gene responses. We found two cell surface receptors likely involved in the detection of *Ilyonectria* and *Stagonosporopsis* taxa and genes in key defense and colonization-response pathways, such as in the modulation of fungal growth, cell wall integrity sensing, Ca^2+^ influx and oxidative burst response, and transcriptional reprogramming of growth and immunity tradeoffs.

In conclusion, we introduce MENTOR as a powerful tool for leveraging multiplex biological networks to identify relationships among genes in a gene set using an intuitive dendrogram visualization. By using this tool to identify the key biological relationships present within a gene set, and distributing this interpretation in a team setting, we hope to streamline the interpretation of omics datasets.

## Acknowledgements/Funding Sources

This manuscript has been authored by UT-Battelle, LLC under Contract No. DE-AC05-00OR22725 with the U.S. Department of Energy. The United States Government retains and the publisher, by accepting the article for publication, acknowledges that the United States Government retains a non-exclusive, paid-up, irrevocable, world-wide license to publish or reproduce the published form of this manuscript, or allow others to do so, for United States Government purposes. The Department of Energy will provide public access to these results of federally sponsored research in accordance with the DOE Public Access Plan (http://energy.gov/downloads/doe-public-access-plan). This work was supported by the Office of Science, Office of Biological and Environmental Research, of the US Department of Energy under Award Numbers DE-AC02-05CH11231, DE-AC02-06CH11357, DE-AC05-00OR22725, and DE-AC02-98CH10886, as part of the DOE Systems Biology Knowledgebase. This research used resources of the Oak Ridge Leadership Computing Facility, which is a DOE Office of Science User Facility supported under contract DE-AC05-00OR22725. Funding was provided by the Plant-Microbe Interfaces (PMI) Scientific Focus Area supported by the Genomic Sciences Program of the Office of Biological and Environmental Research in the DOE Office of Science. The metatranscriptome sequencing conducted by the US Department of Energy Joint Genome Institute is supported by the Office of Science of the US Department of Energy under contract no. DE-AC02-05CH11231. Support for the poplar GWAS SNP dataset is provided by the US Department of Energy, Office of Science Biological and Environmental Research (BER) via the Bioenergy Science Center (BESC) under contract no. DE-PS02-06ER64304. This work was also supported by NIH grants DA051908, DA051913, and DA054071.

## Author Contributions

- Alice Townsend: Methodology; Software; Visualization; Writing - Original Draft; Writing - Review & Editing
- Daniel A. Jacobson: Conceptualization; Methodology; Writing - Reviewing & Editing; Supervision; Project administration; Funding acquisition
- J. Izaak Miller: Conceptualization; Methodology; Software; Formal analysis; Investigation; Resources; Writing - Original Draft; Writing - Review & Editing; Visualization
- Kyle A. Sullivan: Conceptualization; Methodology; Software; Formal analysis; Investigation; Resources; Writing - Original Draft; Writing - Review & Editing; Visualization
- Mallory Morgan: Formal analysis; Investigation; Writing - Original Draft; Writing - Review & Editing; Visualization
- Manesh Shah: Formal analysis; Writing - Review & Editing
- Matthew Lane: Conceptualization; Methodology; Software
- Mikaela Cashman: Conceptualization; Methodology; Software; Writing - Original Draft; Writing - Review & Editing
- Mirko Pavicic: Conceptualization; Software; Visualization; Writing - Original Draft; Writing - Review & Editing

## Code Availability

The open-source MENTOR code is available at https://github.com/Jacobson-CompSysBio/MENTOR.

---

**Supplementary Figure 1.**
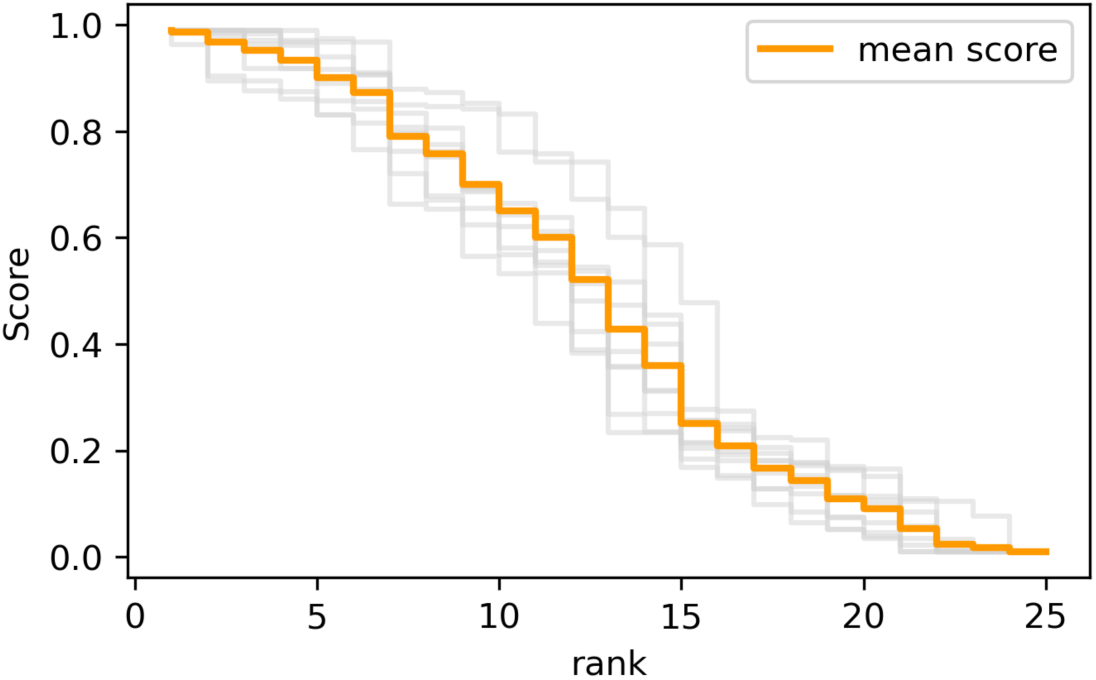
Scores-vs-ranks curve derived from RWR exploration of toy gene set to illustrate elbow point (around score 0.2) at which gene vectors are compared (around top 15 ranks) from the mean score (orange) derived each individual gene used as a seed gene for RWR exploration (grey).

**Supplementary Figure 2.**
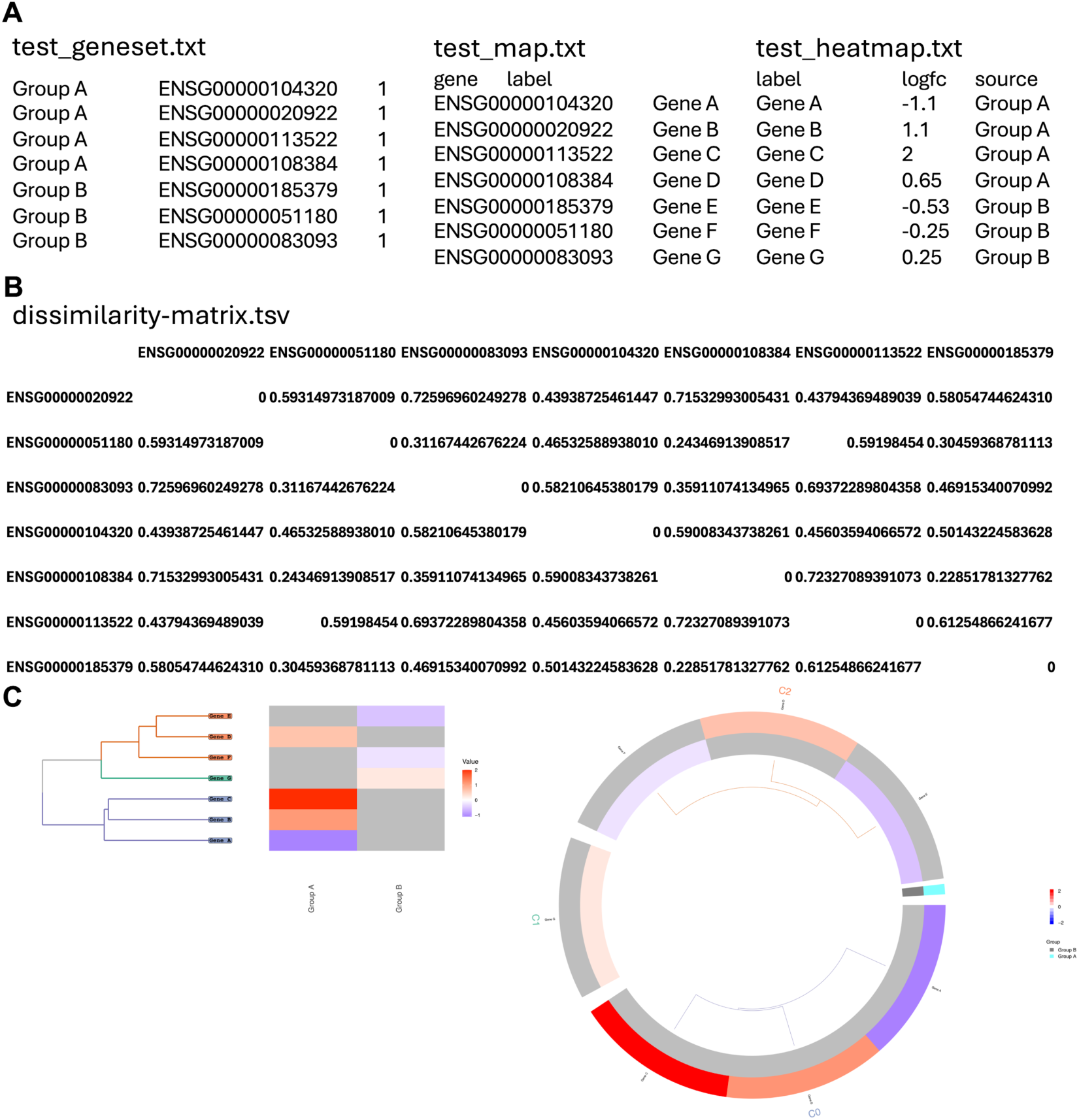
Example inputs and outputs from MENTOR. **A.** Example geneset, map, and heatmap files for MENTOR input. **B.** Example dissimilarity matrix generated as output from MENTOR. **C.** Example of rectangular (left) and circular (i.e. polar, right) dendrogram option from MENTOR output.

**Supplementary Table 1.** GWAS results from *Populus trichocarpa* microbial abundance and labels for MENTOR dendrogram.

