## Supplementary Figures for "MENTOR: Multiplex Embedding of Networks for Team-Based Omics Research"

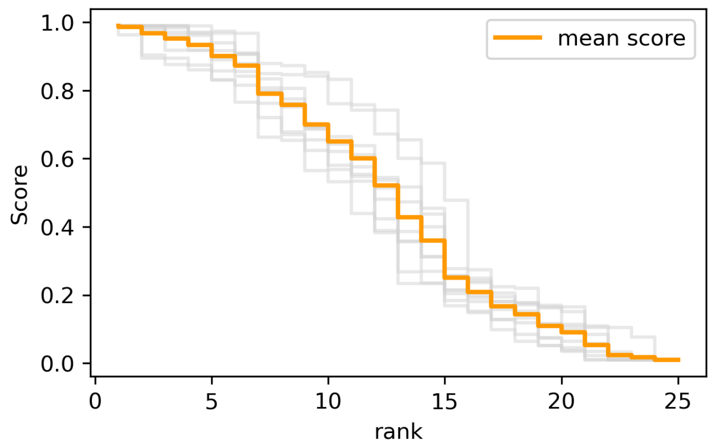


**Supplementary Figure 1.** Scores-vs-ranks curve derived from RWR exploration of toy gene set to illustrate elbow point (around score 0.2) at which gene vectors are compared (around top 15 ranks) from the mean score (orange) derived each individual gene used as a seed gene for RWR exploration (grey).

**A**

test_geneset.txt

| | | |
|---|---|---|
| Group A | ENSG00000104320 | 1 |
| Group A | ENSG00000020922 | 1 |
| Group A | ENSG00000113522 | 1 |
| Group A | ENSG00000108384 | 1 |
| Group B | ENSG00000185379 | 1 |
| Group B | ENSG00000051180 | 1 |
| Group B | ENSG00000083093 | 1 |

test_map.txt

| gene | label |
|---|---|
| ENSG00000104320 | Gene A |
| ENSG00000020922 | Gene B |
| ENSG00000113522 | Gene C |
| ENSG00000108384 | Gene D |
| ENSG00000185379 | Gene E |
| ENSG00000051180 | Gene F |
| ENSG00000083093 | Gene G |

test_heatmap.txt

| label | logfc | source |
|---|---|---|
| Gene A | -1.1 | Group A |
| Gene B | 1.1 | Group A |
| Gene C | 2 | Group A |
| Gene D | 0.65 | Group A |
| Gene E | -0.53 | Group B |
| Gene F | -0.25 | Group B |
| Gene G | 0.25 | Group B |

**B**

dissimilarity-matrix.tsv

| | ENSG0000020922 | ENSG0000051180 | ENSG0000083093 | ENSG0000104320 | ENSG0000108384 | ENSG0000113522 | ENSG0000185379 |
|---|---|---|---|---|---|---|---|
| ENSG0000020922 | 0 | 0.59314973187009 | 0.72596960249278 | 0.43938725461447 | 0.71532993005431 | 0.43794369489039 | 0.58054744624310 |
| ENSG0000051180 | 0.59314973187009 | 0 | 0.31167442676224 | 0.46532588938010 | 0.24346913908517 | 0.59198454 | 0.30459368781113 |
| ENSG0000083093 | 0.72596960249278 | 0.31167442676224 | 0 | 0.58210645380179 | 0.35911074134965 | 0.69372289804358 | 0.46915340070992 |
| ENSG0000104320 | 0.43938725461447 | 0.46532588938010 | 0.58210645380179 | 0 | 0.59008343738261 | 0.45603594066572 | 0.50143224583628 |
| ENSG0000108384 | 0.71532993005431 | 0.24346913908517 | 0.35911074134965 | 0.59008343738261 | 0 | 0.72327089391073 | 0.22851781327762 |
| ENSG0000113522 | 0.43794369489039 | 0.59198454 | 0.69372289804358 | 0.45603594066572 | 0.72327089391073 | 0 | 0.61254866241677 |
| ENSG0000185379 | 0.58054744624310 | 0.30459368781113 | 0.46915340070992 | 0.50143224583628 | 0.22851781327762 | 0.61254866241677 | 0 |

**C**

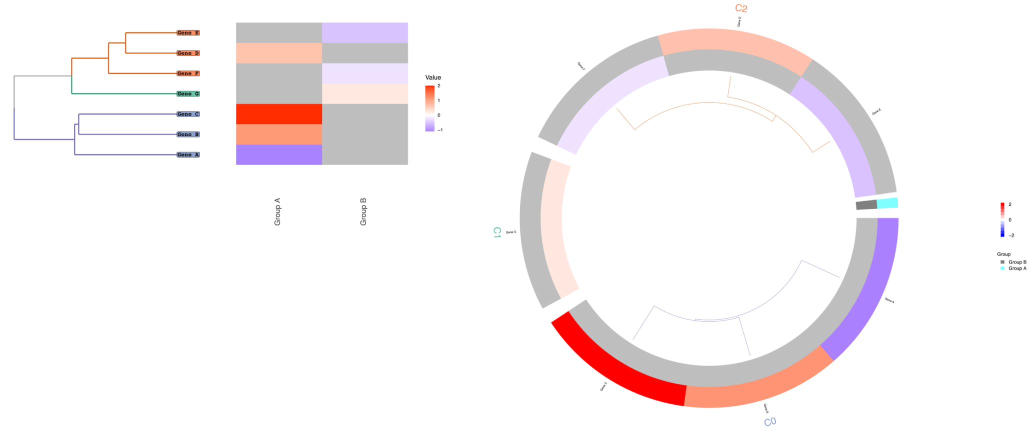


**Supplementary Figure 2. Example inputs and outputs from MENTOR. A.** Example geneset, map, and heatmap files for MENTOR input. **B.** Example dissimilarity matrix generated as output from MENTOR. **C.** Example of rectangular (left) and circular (i.e. polar, right) dendrogram option from MENTOR output.

**Supplementary Table 1.** GWAS results from *Populus trichocarpa* microbial abundance and labels for MENTOR dendrogram.
